# Predicting Empathy from Resting State Brain Connectivity: A Multivariate Approach

**DOI:** 10.1101/539551

**Authors:** Leonardo Christov-Moore, Nicco Reggente, Pamela K. Douglas, Jamie D. Feusner, Marco Iacoboni

## Abstract

Recent task fMRI studies suggest that individual differences in trait empathy and empathic concern are mediated by patterns of interaction between self-other resonance and top-down control networks that are stable across task demands. An untested implication of this hypothesis is that these stable patterns of interaction should be visible even in the absence of empathy tasks. Using machine learning, we demonstrate that patterns of *resting state fMRI connectivity* (i.e. the degree of synchronous BOLD activity across multiple cortical areas in the absence of explicit task demands) of resonance and control networks predict trait empathic concern (n=58). Empathic concern was also predicted by connectivity patterns within the somatomotor network. These findings further support the role of resonance-control network interactions and of somatomotor function in our vicariously-driven concern for others. Furthermore, a practical implication of these results is that it is possible to assess empathic predispositions in individuals without needing to perform conventional empathy assessments.

## 1. Introduction

Empathy is a complex phenomenon that allows us to share in (or *resonate* with) the internal states of others, as well as *infer* their beliefs and intentions (Christov-Moore and Iacoboni, 2016; Decety and Jackson, 2006; Zaki and Ochsner, 2012). It has been suggested that empathy’s *purpose*, in both humans and nonhuman animals, can be broadly divided into two categories: First, promoting pro-social, cooperative behavior via empathic concern for others and second, inferring and predicting the internal states, behavior and intentions of others (Davis, 1983; Preston and De Waal, 2002; Smith, 2006). In this study, we will focus primarily on elucidating the mechanisms underlying empathic concern.

In order to fulfill these purposes, empathy relies in part on our brains’ ability to reflexively process the observed or inferred experiences of others much in the same way we do our own, causing us to respond vicariously to their pain, visceral sensations, and emotions, and simulate their behavior within our own motor systems (reviewed in Zaki and Ochsner, 2012). Furthermore, this phenomenon extends beyond perception to behavior: we tend to reflexively mimic each other’s behavior, often without our knowledge (Chartrand and Bargh, 1999; Lakin and Chartrand, 2003; Meltzoff, 1970; Sperduti et al., 2014), a process that can occur involuntarily when certain prefrontal control areas are damaged (Lhermitte, 1983; De Renzi et al., 1996). We will refer to this reflexive and embodied ability to simulate others as “self-other resonance” (Christov-Moore and Iacoboni, 2016), or *resonance* for short. The most likely neural substrate for resonance appears to be “neural resonance” (Zaki and Ochsner, 2012), the phenomenon of shared neural representations for the perception and experience of disgust (Jabbi et al., 2011; Wicker et al., 2003), somatosensation (Bufalari et al., 2007; Masten et al., 2011; Singer et al., 2006), emotion (Carr et al., 2003; Pfeifer et al., 2008) and motor behavior (Iacoboni, 2009; Keysers and Fadiga, 2008). Not surprisingly, neural resonance has been repeatedly associated with self-reported measures of trait empathy (Avenanti et al., 2009; Jabbi et al., 2011; Pfeifer et al., 2008) and is predictive of pro-social behavior (non-strategic generosity in economic games: Christov-Moore and Iacoboni, 2016; harm aversion in moral dilemmas: Christov-Moore et al., 2017b; donations to reduce pain in another: Gallo et al., 2018; helping behavior: Hein et al., 2011; Morelli et al., 2014; charitable donations: Ma et al., 2011), suggesting that our resonance with others may underlie our *empathic concern* (and hence prosocial inclinations) for others.

In further support of a common substrate, prosocial inclinations and self-other resonance are similarly modulated by others’ closeness, status, group affiliation and perceived trustworthiness (Chartrand and Bargh, 1999; Cheng et al., 2010; Gu and Han, 2007; Guo et al., 2012; Hein and Singer, 2008; Lakin and Chartrand, 2003; Lamm et al., 2007; Liew et al., 2007; Loggia et al., 2008; Reynolds-Losin et al., 2013; 2014; Schmälzle et al., 2017; Singer et al., 2006; Sperduti et al., 2014). This is likely due to top-down control processes that integrate contextual information and conscious appraisal with affective, somatosensory and motor processes into behavior and decision-making, implemented by prefrontal and temporal systems including the temporoparietal junction (TPJ) as well as dorsomedial and dorsolateral prefrontal cortex (DMPFC and DLPFC) (Banks et al., 2007; Brighina et al., 2010; Cho and Strafella, 2009; Decety and Lamm, 2007; Miller and Cohen, 2001; Spengler et al., 2010; Tassy et al., 2012; Volman et al., 2011; Winecoff et al., 2013). Not surprisingly, these control systems overlap considerably with those associated with conscious appraisal processes and inferential forms of empathy or mentalizing (Mahy et al., 2014). The nature of this control seems to be inhibitory: a recent study has found that disruptive neuromodulation of DMPFC and DLPFC caused a decrease in the inhibitory influence of context on prosocial behavior (Christov-Moore et al., 2017a). Evidence suggests that this top-down control of resonance is also continuously engaged: lesions to prefrontal cortex are associated with compulsive imitative behavior, suggesting that, for normal behavior to exist, some mechanisms to control resonance are always at play, unless damaged (Lhermitte, 1983; De Renzi et al., 1996). Within the context of empathy, resonance and control may exist most often as clusters within a single integrated system.

Indeed, the neural bases of resonance and control processes are not cleanly separable within cognitive function. Recent research suggests that somatomotor and affective processing contribute to our evaluations of others’ internal states, beliefs, and intentions (Christov-Moore and Iacoboni, 2016; Christov-Moore et al., 2017a; Frith and Singer, 2008; Gallese, 2007; Obhi, 2012; Schulte-Ruther et al., 2007), as well as our decisions about others’ welfare (Camerer, 2003; Christov-Moore et al., 2017a, 2017b; Hewig et al., 2011; Greene et al., 2001; Oullier and Basso, 2010; Van ’t Wout et al., 2006). Conversely, top-down control processes are increasingly implicated in the contextual modulation of neural resonance (Cheng et al., 2010; Gu and Han, 2007; Guo et al., 2012; Hein and Singer, 2008; Lamm et al., 2007; Loggia et al., 2008; Reynolds-Losin et al., 2013; 2014; Singer et al., 2006). Many studies have reported concurrent activation of and connectivity between networks associated with resonance and top-down control, such as during passive observation of emotions or pain (Christov-Moore and Iacoboni, 2016), passive observation of films depicting personal loss (Raz et al., 2014), reciprocal imitation (Sperduti et al., 2014), tests of empathic accuracy (Zaki and Ochsner, 2009), and comprehension of others’ emotions (Spunt and Lieberman, 2012). Co-existence of bottom-up resonance and top-down control mechanisms can be documented even at the level of TMS-induced motor evoked potentials (MEPs), a functional readout of motor excitability (Gordon et al, 2018). Thus, the neural instantiation of resonance and control may rely on systems that operate like connected clusters in a network, with different modes and configurations of function (Fox and Friston, 2012).

On the basis of this evidence, we propose that individual differences in empathic function (*particularly empathic concern for others*) arise in large part from stable, characteristic interactions between resonance and control processes at the neural level (Christov-Moore and Iacoboni, 2016; Christov-Moore et al., 2017a). This view is in line with studies showing that individual differences in active, task-relevant network configuration are reflected in intrinsic functional connectivity patterns (Smith et al., 2009; Tavor et al., 2016). We propose that in adults, these individual differences in empathic function should be apparent in resting connectivity (i.e. in the absence of empathy-evoking stimuli), much in the way a river carves out a characteristic pattern in bedrock over time. If so, this could have implications for understanding differences in empathic functioning without needing to probe participants with specialized tasks or questionnaires. Thus, we approached this current work with specific and general hypotheses: Specifically, we hypothesized, in line with our prior studies on the neural bases of prosociality, that multivariate characteristics of the interaction between resonance and top-down control networks (proposed in Christov-Moore and Iacoboni, 2016) would predict participants’ empathic concern for others. In contrast to the previous univariate analyses, our goal was to derive a multivariate understanding of how empathy is represented by connectivity in these networks. Following work on resting-state and empathy (e.g. Cox et al., 2012), in a more exploratory fashion, we hypothesized that resting connectivity patterns within and between other cortical networks could also predict levels of trait empathy, with particular attention to the somatomotor network, which has been linked to many forms of prosociality (non-strategic generosity in economic games: Christov-Moore and Iacoboni, 2016; harm aversion in moral dilemmas: Christov-Moore et al., 2017b; donations to reduce pain in another: Gallo et al., 2018; helping behavior: Hein et al., 2011; Morelli et al., 2014; charitable donations: Ma et al., 2011).

Additionally, there is a great deal of evidence for sex differences in empathy across a broad array of measures and associated brain function (Cohn, 1991; Christov-Moore et al., 2014; Eisenberg and Lennon, 1983; Feingold, 1994; Hall, 1978; 1984; Hoffman, 1977; Konrath et al., 2011; Thompson and Voyer, 2014; although, for negative/null results see Lamm et al., 2011). For example, females display greater concern and sympathetic behavior towards others in real and hypothetical scenarios (Christov-Moore et al., 2014; Eisenberg and Lennon 1993; Friesdorf et al., 2015; Mesch et al. 2011). Females also show greater vicarious somatosensory responses to the sight or knowledge of another person in pain or distress (Christov-Moore et al., 2014; Christov-Moore and Iacoboni, 2018; Groen et al., 2013; Singer et al., 2006; Yang et al. 2009), and exhibit greater facial mimicry when viewing emotional facial expressions (Sonnby-Borgström 2002). For this reason, we controlled for sex within the primary analysis predicting trait empathy.

Taken together, previous studies suggest that resonance and control processes’ interactions may be the basis for individual differences in empathic concern for others, and that these interactions are relatively stable across task demands, Thus, we sought to test two families of hypotheses: I) our primary, theory-driven hypothesis that Resonance and Control interconnectivity at rest predicts trait Empathic Concern, and II) our exploratory, theory-consistent but broader hypothesis that trait empathy can be predicted from resting intra- and inter-connectivity of intrinsic brain networks.

## 2. Materials and methods

### 2.1. Participants

Participants were 58 ethnically diverse adults aged 18-35 (30 female, 28 males) recruited from the local community through fliers. All recruitment and experimental procedures were performed under approval of UCLA’s institutional review board, in accordance with the ethical standards of the institutional and/or national research committee and with the 1964 Helsinki declaration and its later amendments or comparable ethical standards. Informed consent was obtained from all individual participants included in the study.

Eligibility criteria for participants included: right handed, no prior or concurrent diagnosis of any neurological, psychiatric, or developmental disorders, and no history of drug or alcohol abuse. These were all assessed during preliminary screening interviews conducted by phone at the time of recruitment.

### 2.2. Trait empathy assessment

Participants filled out the *Interpersonal Reactivity Index* (IRI) at the end of each experimental session in a closed room, unobserved. The IRI (Davis, 1983) is a widely used (Avenanti et al., 2009, Pfeifer et al., 2008) and validated (Litvack-Miller et al., 1997) questionnaire designed to measure both “cognitive” and “emotional” components of empathy. It consists of 24 statements that the participant rates on a 5-point scale ranging from 0 (Does not describe me very well) to 5 (Describes me very well). The statements are calculated to test four theorized subdimensions of empathy:

*Fantasizing Scale (FS)*: the tendency to take the perspective of fictional characters.
*Empathic Concern (EC)*: sympathetic reactions to the distress of others.
*Perspective Taking (PT)*: the tendency to take other’s perspective.
*Personal Distress (PD)*: aversive reactions to the distress of others

Participants’ scores were summed for each sub-dimension (measured by 6 items) to make 4 scores per participant. Cronbach’s alpha, a measure of reliability, was assessed for the IRI using SPSS (FS=0.752, EC=0.792, PT=0.816, PD=0.839).

### 2.3. Functional MRI data collection

All neuroimaging data was acquired via a series of MRI scans conducted in a Siemens Trio 3T scanner housed in the Ahmanson-Lovelace Brain Mapping Center at UCLA. Resting data were collected while participants passively observed a white fixation cross on a black screen. They were instructed only to “Look at the fixation cross and just let your mind wander.” Resting-state functional images were acquired over 36 axial slices covering the whole cerebral volume using an echo planar T2*-weighted gradient echo sequence (6 minutes; TR=2500 ms; TE=25 ms; flip angle=90 degrees; matrix size=64 x 64; FOV 20 cm; in-plane resolution=3 mm x 3 mm; slice thickness=3 mm/1 mm gap). A T1-weighted volume was also acquired in each participant (TR=2300 ms, TE=25 ms, TI=100 ms, flip angle=8°, matrix size=192×192, FOV=256 cm, 160 slices, voxel size 1.3 x 1.3x 1.0 mm).

### 2.4. Functional MRI preprocessing

Functional MRI preprocessing was performed in FEAT (FMRI Expert Analysis Tool), part of FSL (FMRIB’s Software Library, www.fmrib.ox.ac.uk/fsl). After motion correction using MCFLIRT, images were temporally high-pass filtered with a cutoff period of 100 seconds (equivalent to .01Hz) and smoothed using a 6 mm Gaussian FHWM algorithm in 3 dimensions. Our protocol stipulated that participants showing absolute or relative head motion exceeding 1mm were excluded from further analyses, though no participants exceeded this threshold. In order to remove non-neuronal sources of coherent oscillation in the relevant frequency band (.01-.1Hz), preprocessed data was subjected to probabilistic independent component analysis as implemented in MELODIC (Multivariate Exploratory Linear Decomposition into Independent Components) Version 3.10, part of FSL (FMRIB’s Software Library, www.fmrib.ox.ac.uk/fsl). Noise components corresponding to head motion, scanner noise, cardiac/respiratory signals were identified by observing their localization, time series and spectral properties (as per Kelly et al., 2010) and removed using FSL’s regfilt command. Each participants’ functional data was coregistered to standard space (MNI 152 template) via registration of an averaged functional image to the high resolution T1-weighted volume using a six degree-of-freedom linear registration and of the high-resolution T1-weighted volume to the MNI 152 template via nonlinear registration, implemented in FNIRT.

### 2.5. Designation of regions of interest

All networks were created by pooling from a set of 198 5mm spherical ROI’s. 196 of the ROIs were derived from a functionally-derived cortical atlas (Power et al. 2011). We also included an additional pair of 5mm ROI’s centered on left (x=-22mm, y=-6mm, z=-14mm) and right (x=22mm, y=-6mm, z=-14mm) amygdala, as this region was not included in the original cortical atlas. We used ROIs from the following networks defined by Power et al. (2011): Visual (31 ROIs), Fronto-parietal (25 ROIs), Somatosensory-Motor (25 ROIs), Dorsal Attention (11 ROIs), Ventral Attention (9 ROIs), Salience (18 ROIs), Memory Retrieval (5 ROIs), Cingulo-Opercular (14 ROIs), and Default Mode (58 ROIs) Networks. ROIs were defined in MNI_152 standard space.

Two theory-driven networks (bottom up resonance and top down control) were also created by selecting ROIS from the Power cortical atlas overlapping with brain areas associated with neural resonance and top-down control. Resonance areas included the core cortical imitation circuitry (inferior frontal gyrus, inferior parietal lobule, superior temporal sulcus), as well as insular, limbic (bilateral amygdala) and somatomotor areas associated with neural resonance for visceral sensation, emotion, pain and motor behavior (e.g. reviewed in Lamm, 2011, Zaki & Ochsner, 2012). This putative bottom up resonance network consisted of 34 ROIs. Control areas included dorsolateral prefrontal cortex, temporoparietal junction, lateral orbitofrontal cortex and sites covering a range from dorsal to ventral medial prefrontal and paracingulate cortex, implicated in top-down regulation of spontaneous and deliberate imitation, affect and pain (Banks et al., 2007; Brighina et al., 2010; Cho and Strafella, 2009; Christov-Moore and Iacoboni, 2016; Christov-Moore et al., 2017b; Decety and Lamm, 2007; Miller and Cohen, 2001; Spengler et al., 2010; Tassy et al., 2012; Volman et al., 2011; Winecoff et al., 2013). This putative top-down control network consisted of 22 ROIs. This allowed us to test our conceptual model of resonance-control interaction as a substrate for empathic concern (Christov-Moore & Iacoboni, 2016, Christov-Moore et al., 2017b), while constraining ROI locations to those defined in Power et al. (2011) and assigning these ROI locations to the two networks on the basis of existing literature. See Table 1 for a list of ROIs used to define the resonance and control networks and Figure 1 for a visual rendering of the same ROIs/networks.

**Figure 1.**
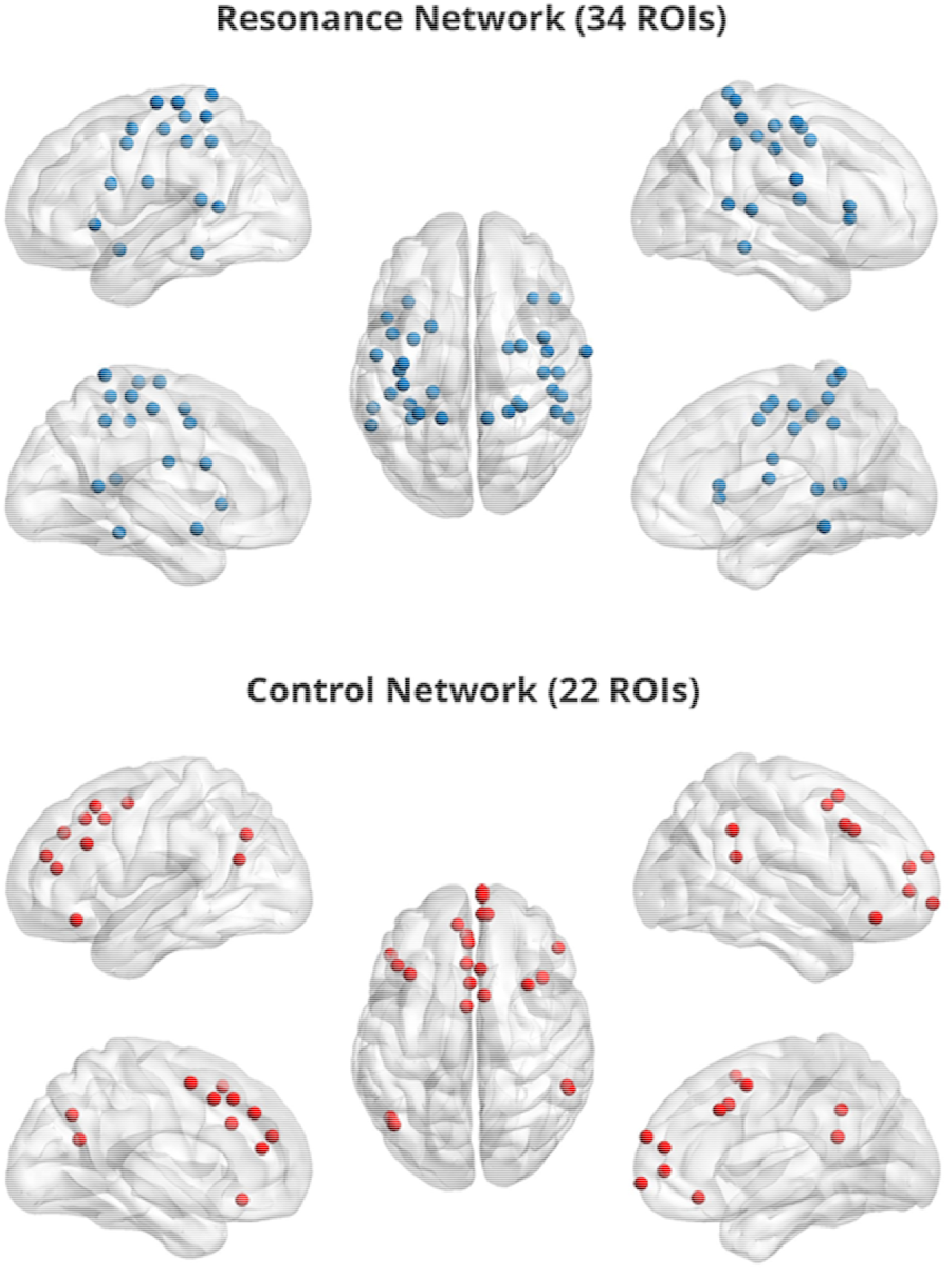
Resonance (top) and Control (bottom) Networks. 5mm regions of interest were visualized with the BrainNet Viewer (http://www.nitrc.org/projects/bnv/) (Xia et al., 2013).

**Table 1.**
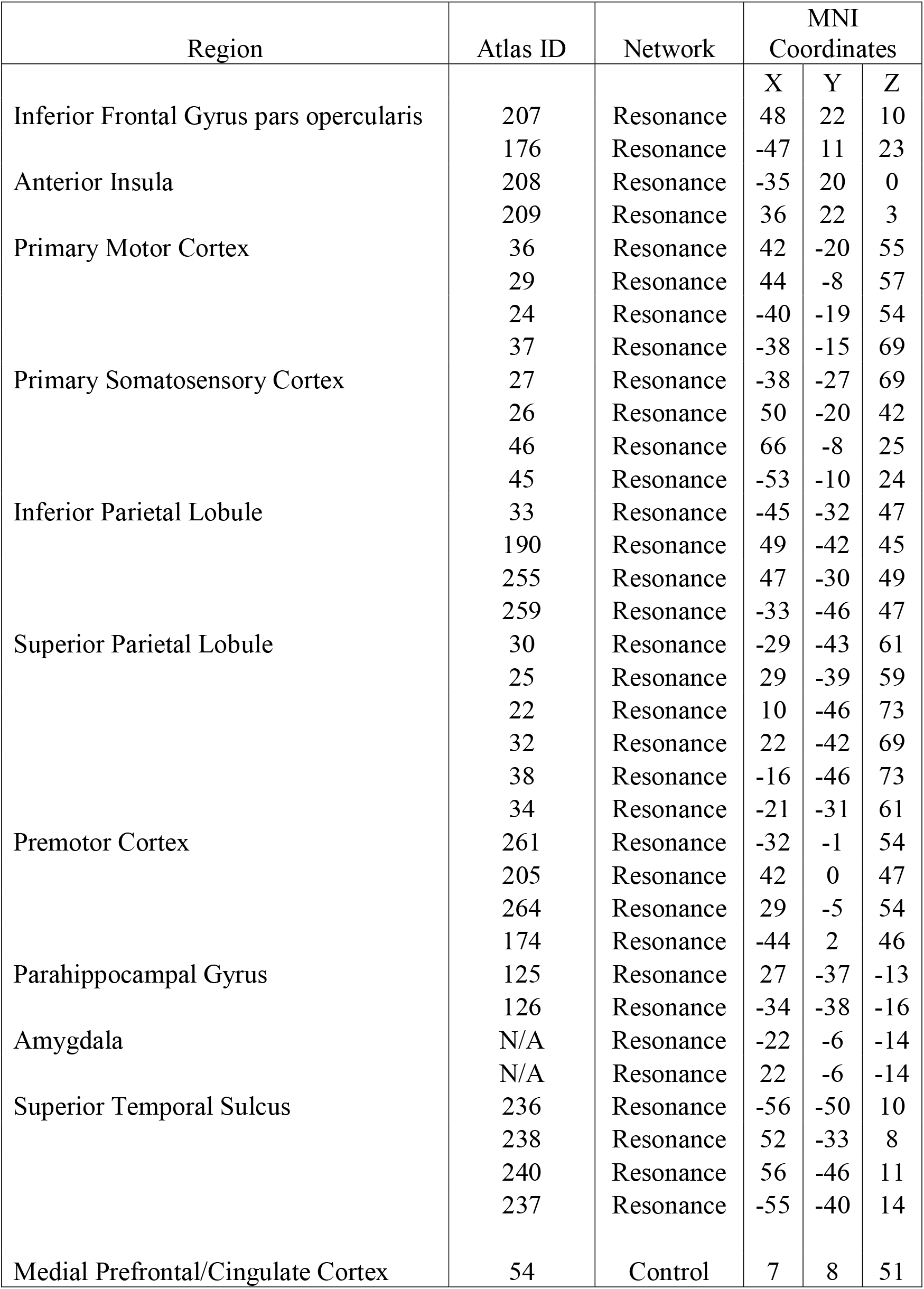
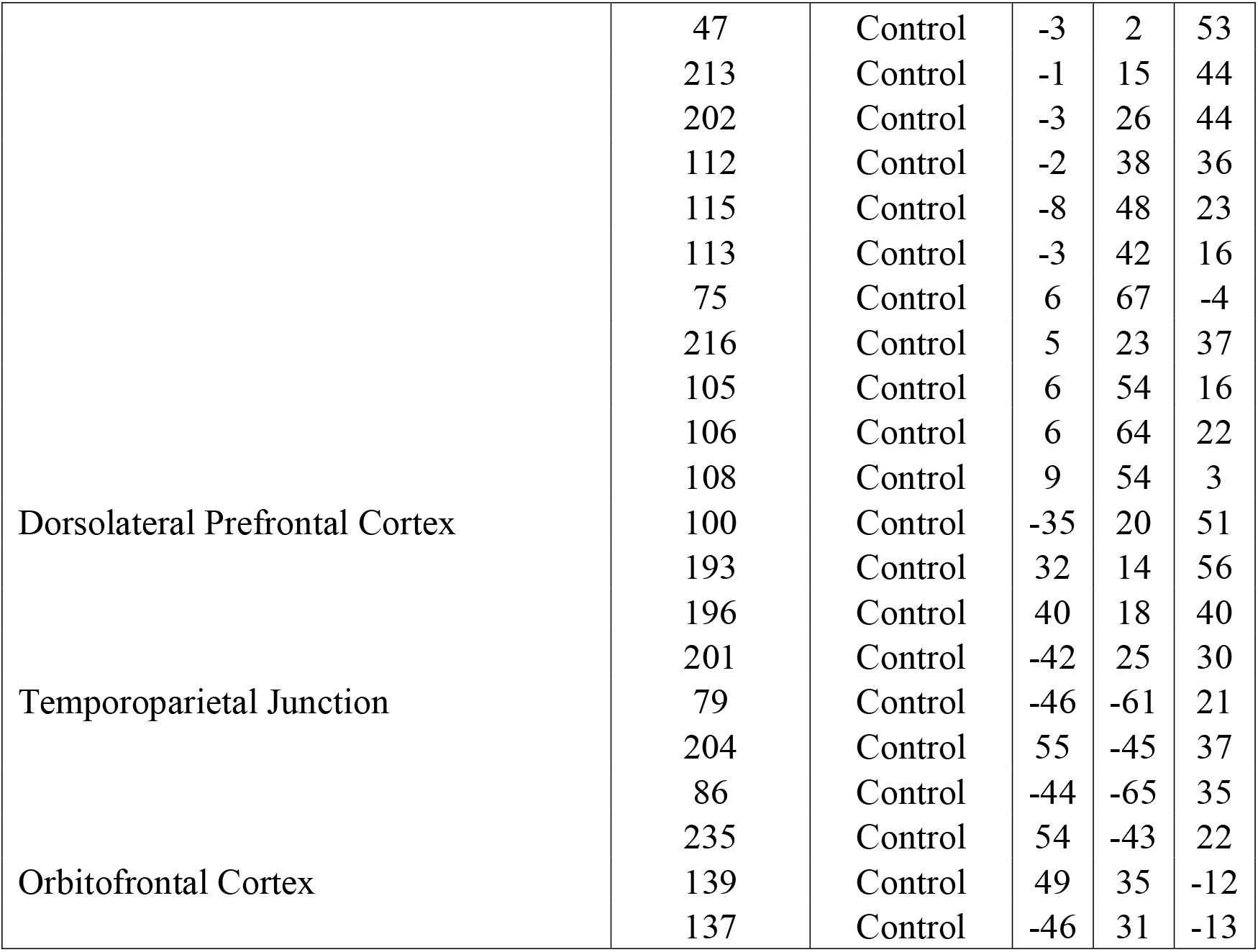
MNI coordinates of Powers cortical atlas ROI’s employed in resonance and control networks. IFGpo= Inferior Frontal Gyrus pars opercularis

### 2.6. Machine learning analyses

Mean BOLD time-courses were extracted from the average activity across voxels within each ROI. Matrices of pairwise Pearson correlation coefficients were created for each participant by correlating each ROI’s BOLD time-course with every other ROI within each network, creating a set of features (non-redundant functional connectivity values) for each participant and network. For “between-networks” analyses, ROIs belonging to each pair of networks being studied (e.g. Resonance and Control) were combined into a single network, allowing for pairwise connectivity across all member ROIs.

To account for potential covariation, participant sex was iteratively regressed out of each feature and the residuals were subsequently used as the functional connectivity features. We implemented a leave-ten-subjects-out cross validation to assess the predictive power of network-specific feature sets. Specifically, we leveraged a least absolute shrinkage and selection operator (LASSO) regression model built on N-10 participants’ feature sets for each IRI subscale. The model’s intercept term and outcome beta values were then used as coefficients for each left-out subject’s feature set—obtaining a predicted subscale measure for that individual. After N folds, whereby each set of 10 participants was left out exactly once, we correlated the array of predicted values (*Ŷ*) with the actual values (Y), yielding Pearson’s R– a measure of our model’s feature-dependent ability to capture the behavioral variance across participants. We repeated this cross-validation 10 times and averaged the R values to converge on a true estimate of our test statistic, independent of which participants were randomly included in each fold. The LASSO regularization parameter was optimized before the leave-ten-subject-out cross-validation by using the least angle regression (LARS) algorithm on an N - 1 cross-validation that maximized the Pearson correlation between predicted values (Ӯ) with the actual values (Y).

### 2.7. Significance testing and multiple comparisons correction

R-values from the N-10 cross-validation, averaged across the ten iterations, were submitted to a significance test of the correlation coefficient 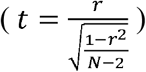. In order to correct for multiple comparisons, we applied three family-wise corrections, for each set of hypotheses: I) our main, theory-driven hypothesis that Resonance and Control interconnectivity predict Empathic Concern) and II) our exploratory, broad hypothesis that trait empathy can be predicted from resting intra- and inter-connectivity of canonical and theory-driven intrinsic networks. Matrices of p-values within each family were created using a Benjamini-Hochberg approach (Benjamini and Hochberg, 1995) in R (p.adjust; method=“BH”; R Core Team, 2013) and corrected p-values were considered significant at the 5% positive-tail (i.e. p<0.05). (Negative R values, indicating poor prediction accuracy – i.e. predicting a negative subscale score when the actual value is positive – are not readily interpretable.)

### 2.8. Data and code availability statement

All data are freely available upon request. For human fMRI and behavioral data contact Leonardo Christov-Moore. Custom scripts used in this analysis can be found at https://github.com/mobiuscydonia/Moore-2019-Empathic-Concern. Data and code sharing adopted by the authors complies with the requirements of our funding body as well as institutional ethics.

## 3. Results

### 3.1. Trait empathy IRI scores

**Table 2.** Means (with 95% confidence intervals) and standard deviations for each IRI subscale by gender. Females scored higher on personal distress (PD), but not on PT (perspective-taking), FS (fantasizing) or EC (empathic concern).

| | $\bar{x}_{\square}$ (95%CI) | $\sigma_{x_{\square}}$ | $\bar{x}_{\square}$ (95%CI) | $\sigma_{x_{\square}}$ |
| --- | --- | --- | --- | --- |
| FS | 18.58(16.54,20.61) | 5.02 | 20.84(18.89,22.79) | 4.82 |
| EC | 22.61(20.88,24.36) | 4.31 | 24.54(22.79,26.29) | 4.34 |
| PT | 20.00(17.88,22.12) | 5.24 | 20.73(18.78,22.68) | 4.82 |
| PD | 11.73(9.41,14.05) | 5.75 | 16.54(14.39,18.69) | 5.32 |

We used a one-way ANOVA to examine whether the male and female participants differed significantly in self-reported trait empathy. Males and females did not differ significantly in Fantasizing (F=2.68, p=.108), Empathic Concern (F=2.59, p=.114), or Perspective-Taking (F=.274, p=.603). However, female subjects scored significantly higher on Personal Distress (F=9.79, p=.003).

### 3.2. Machine learning and connectivity

As described above, for these analyses, we examined 5mm spherical regions of interest set in MNI_152 space for the Visual, Fronto-Parietal, Cingulo-Opercular, Dorsal and Ventral Attention, Salience, Memory Retrieval, Subcortical, Somatomotor, and Default Mode Networks (derived from Power et al., 2011) as well as two theory-driven networks (Resonance and Control, see figure 1) created based on a) a model of resonance-control interactions as a substrate for empathic concern (Christov-Moore and Iacoboni, 2016) and with ROIs derived from the literature as described above.

#### 3.2.1. Within-network resting connectivity predicts trait empathy

When examining the predictive power of connectivity weights *within* the selected intrinsic networks (Figure 2), empathic concern was significantly predicted by the somatomotor network (R=0.374, p=0.022, Benjamini-Hochberg false discovery rate – FDR – corrected). Personal distress was predicted above threshold by resonance (R=0.236, p=0.037, uncorrected), control (R=0.22, p=0.048, uncorrected) and cingulo-opercular networks (R=0.242, p=0.033, uncorrected), however these did not survive FDR correction for multiple comparisons.

**Figure 2.**
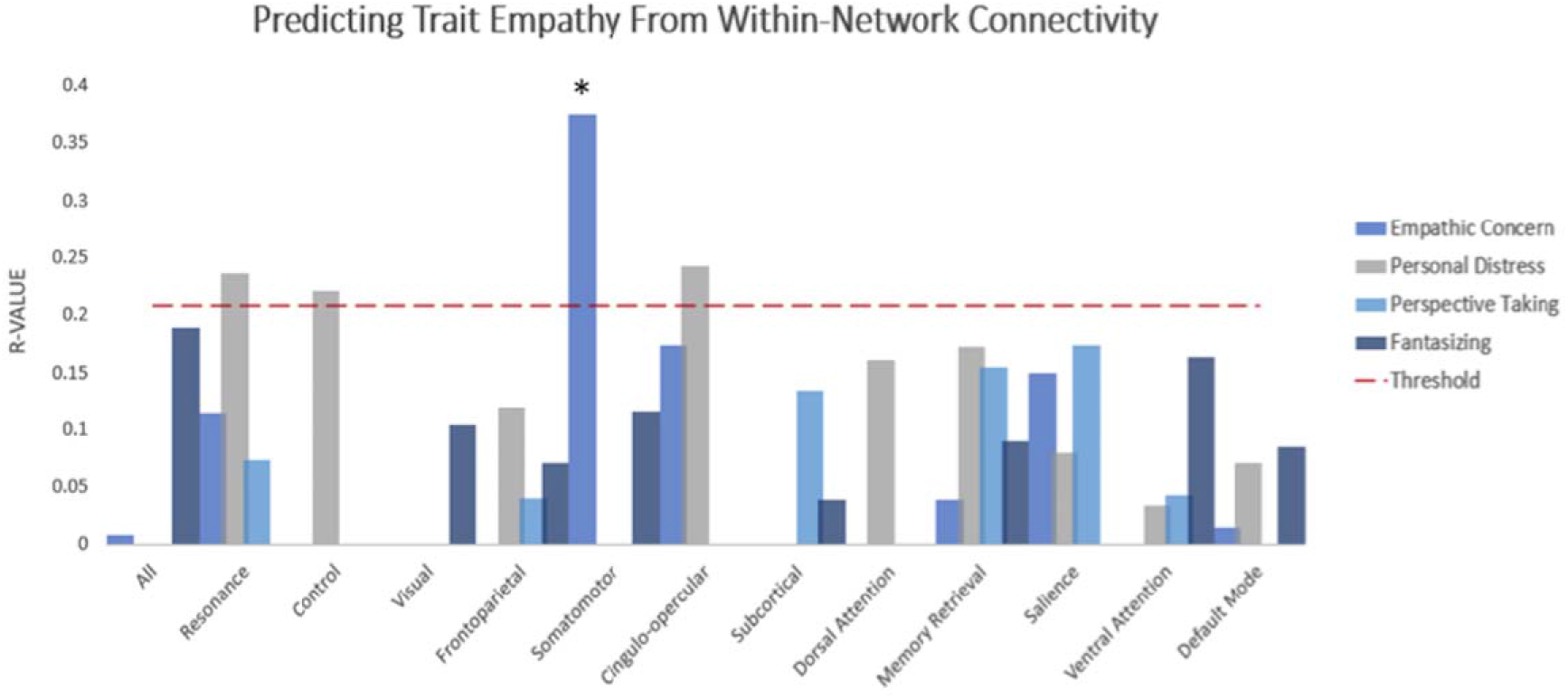
Within-Network Somatomotor Resting Connectivity Predicts Empathic Concern. Y-axis depicts average correlations between values predicted from model trained on n-10 cross-validation set and remaining 10 subjects over multiple iterations. Red dashed bar indicates threshold for p<0.05, uncorrected. * p-value <0.05 FDR corrected.

#### 3.2.2. Between-network resting connectivity predicts trait empathy

When analyzing the predictive power of interactions within multiple networks simultaneously, empathic concern was predicted by the interaction between the a priori resonance and control networks (R=0.221, p=0.0475, Benjamini-Hochberg false discovery rate – FDR – corrected), supporting the primary hypothesis of this study.

When testing our second family of hypotheses, we examined three types of between-network complexes: bottom-up resonance (visual/somatomotor, visual/frontoparietal, somatomotor/frontoparietal), resonance and control (control/frontoparietal, control/visual, control/somatomotor, cingulo-opercular/default mode), and links of no a priori interest as a comparison (dorsal/ventral attention, salience/dorsal attention), selected to test whether empathic function could be predicted by differences in attentional networks (Figure 3). None of these survived FDR correction for multiple comparisons

**Figure 3.**
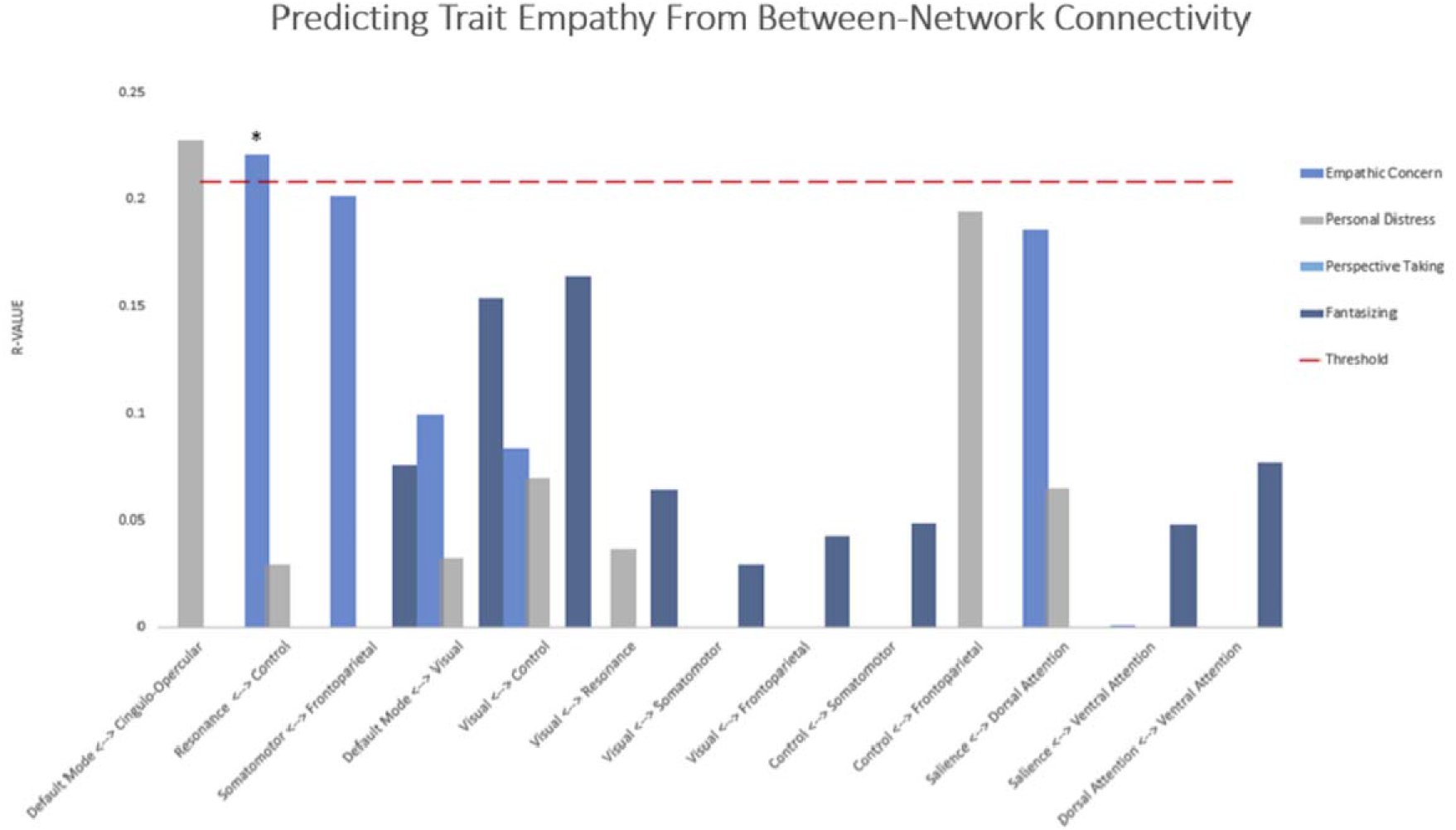
Between-Network Resting Connectivity of Resonance and Control Networks Predicts Empathic Concern. Y-axis depicts average correlations between values predicted from model trained on n-10 cross-validation set and remaining 10 subjects over multiple iterations. Red dashed bar indicates threshold for p<0.05, uncorrected. * p-value <0.05 FDR corrected.

## 4. Discussion

In this study, we tested two hypotheses:

I. We hypothesized that participants’ empathic concern for others would be predicted by resting connectivity between our theory-driven and literature-derived resonance and control networks.
II. We hypothesized that we could predict subcomponents of participants’ trait empathy from the within- and between-network resting connectivity of canonical resting state networks.

As hypothesized in (I), participants’ levels of empathic concern were predicted by patterns of interconnectivity between the resonance and control networks, supporting the hypothesis (put forth in Christov-Moore and Iacoboni, 2016 and supported by Christov-Moore et al., 2017a) that these systems a) continuously interact in a characteristic fashion observable in the absence of pertinent task demands, and b) this interaction is a likely neural substrate of empathic concern for others. Our findings (along with the previous work that prompted this study) support a dynamic, integrated view of empathic function, based on complex patterns of interaction between resonance and control systems rather than simply a univariate measure of overall connectivity. Indeed, numerous studies have reported interactions between resonance and control networks during passive observation of emotions or pain (Christov-Moore and Iacoboni, 2016), reciprocal imitation (Sperduti et al., 2014), tests of empathic accuracy (Zaki and Ochsner, 2009), and comprehension of others’ emotions (Spunt and Lieberman, 2012). Interestingly, Raz et al. (2014) found evidence for complex, context dependent interactions between “simulation” and “theory-of-mind” networks (largely corresponding to what are defined here as resonance and control networks) during empathic experience (observing films depicting personal loss). This multivariate approach may help reconcile findings supporting an integrated view with activation (e.g. Van Overwalle and Baetens, 2009) or lesion studies that suggest dissociated systems (e.g. Shamay-Tsoory, et al., 2009): Lesions (transient/induced or physical) may simply be altering a crucial node for a specific integrated network outcome, just as a hand injury may affect the ability to catch a ball more than a back injury, though catching-like activities typically rely on hands, arms, and the core operating in unison. Indeed, the complexity of these interactions may be an obstacle to their efficient detection by standard activation or univariate connectivity methods. By employing flexible machine learning methods that make few a priori assumptions about the patterns of intrinsic connectivity underlying individual differences, we may achieve a more comprehensive multivariate view of the possible network-level patterns of neural interaction that give rise to individual differences in empathic function. It is common within cognitive neuroscience to theorize first about psychological processes and then investigate the neural correlates of such processes. However, in an exceedingly complex system such as the brain, much could be gained by approaching the problem from the opposite direction, by investigating how psychological processes *emerge* from brain organization (Fox and Friston, 2012).

As for (II), empathic concern was predicted by the within-network connectivity of the somatomotor network. This result further supports an embodied, somatomotor foundation for our concern for others’ welfare, in line with numerous findings relating vicarious somatosensory activation to multiple forms of prosocial behavior ((non-strategic generosity in economic games: Christov-Moore and Iacoboni, 2016; harm aversion in moral dilemmas: Christov-Moore et al., 2017b; donations to reduce pain in another: Gallo et al., 2018; helping behavior: Hein et al., 2011; Morelli et al., 2014; charitable donations: Ma et al., 2011). This also agrees with our recent finding that inferior premotor activation during observation of pain in others was predictive of participants’ later tendency to avoid inflicting harm in hypothetical moral dilemmas (Christov-Moore et al., 2017b). A major proposed subcomponent of empathy is fantasizing (Clay & Iacoboni, 2011, Davis, 1983), our ability to take the perspective of absent or fictional characters and become correspondingly invested in their welfare. Perhaps we implicitly construct internal models of others (present or implied/hypothetical) using perceptual, affective and motor experiences we associate with past experience, framed by others’ intentions, moral character, group affiliation, etc. This embodied model of the “other” and its contextual framing would likely be represented by interactions between resonance and control processes, thus shaping the relative utility of their welfare (Bechara & Damasio, 2005), and hence the positive and negative reward values assigned to the outcomes of decisions that can affect them (Fehr & Camerer, 2007).

A clinical avenue suggested by this study is the potential ability to predict empathic functioning in populations that might have difficulty performing empathy tasks or filling out questionnaires, either due to being less cooperative or less cognitively able, e.g., in populations such as those with schizophrenia, low functioning autism, intellectual disabilities, or traumatic brain injury. Individuals in these groups might have, in principle, intact inherent capability for normal-range empathy that could be impeded by other limitations such as verbal or nonverbal communication (autism) or disorganized thought processes (schizophrenia); thus it would help us know what reasonable outcomes in terms of social and interpersonal functioning could be expected to result from therapies that help with training to rehabilitate or improve empathy, ultimately in the interest of enhancing social competence and social cognition. Indeed, it may be pertinent to include measures of empathic function along with standardized, multisite resting state scan protocols (like the Human Connectome Project), paving the way for a massive data-driven approach to produce models that can predict empathic function from the resting brain in many different populations.

### 4.1. Limitations

While we have focused primarily on network interactions underlying empathic concern, future work could make a similar theory-driven test of putative networks underlying other facets of empathic function (such as perspective-taking). Additionally, while we have shown that network properties, i.e. the aggregate of connectivity weights, can be used meaningfully as features to predict trait empathy, the nature of this multivariate approach does not readily provide simple conclusions about *what aspects of these networks* are predictive and in which direction. We cannot, for example, say: “increased interconnectivity predicts personal distress.” Graph theoretical analyses may allow for complementary mechanistic insights into the properties of whole networks, and parts of networks, that can predict trait empathy. Also, this study only examines “standard” connectivity, i.e. BOLD time-series correlation. Effective connectivity or mutual information analyses may shed light on more complex or nonlinear interactions that might underlie the more dynamic, cognitive aspects of empathy (such as mentalizing or perspective-taking). Further, future larger studies could employ a whole-brain search that could potentially more broadly identify additional systems that contribute to empathy outside of the chosen networks implicated from previous studies.

### 4.2. Conclusions

In conclusion, these findings support a dynamic, integrated model of the neural substrates for empathic concern. The presence of informative patterns of connectivity at rest suggests that these networks interact in a characteristic function regardless of task demands. Along the same lines, albeit at a more fine-grained, local level, these data support an embodied view of empathic concern, in which somatomotor representations of others’ harm situates our feelings and decisions about their welfare.

More broadly, these results add an important piece to a growing body of work demonstrating links between resting and task-positive brain function (Smith et al., 2009), suggesting that the two may not be as cleanly separable as is often implicitly assumed. Perhaps a multivariate, theory-driven method like the one employed here, combined with large datasets, could be applied to predict many aspects of cognition and behavior from resting brain activity. Having metrics that are stable, relatively context-invariant, and predictive of behavior is of great importance for the future of psychiatric research. Along these lines, finding markers of empathic functioning that are visible at rest may be of great potential prognostic and therapeutic utility, and could shed light on mechanisms underlying both healthy and abnormal empathic functioning.

## 5. Acknowledgments

This work was supported by the National Institute of Mental Health under grant R21 MH097178 to M.I., and by the National Science Foundation under a Graduate Fellowship Grant DGE-1144087 to L.C.M. For generous support the authors also wish to thank the Brain Mapping Medical Research Organization, Brain Mapping Support Foundation, Pierson-Lovelace Foundation, The Ahmanson Foundation, William M. and Linda R. Dietel Philanthropic Fund at the Northern Piedmont Community Foundation, Tamkin Foundation, Jennifer Jones-Simon Foundation, Capital Group Companies Charitable Foundation, Robson Family and Northstar Fund. An earlier version of this manuscript (identical in methods and results] has been released as a Pre-Print at BioRxiV (Christov-Moore et al., 2019].

## 7. Contribution to the field statement

Empathy is a cornerstone of human social cognition and prosocial behavior. It aids us in understanding others’ internal states and permits us to share in those states, motivating us to cooperate with and aid one another. Here we aimed to elucidate the neural basis of this *empathic concern*. Modern accounts of brain function increasingly view the brain as a massively interconnected system in which cognitive functions emerge from the interaction between clusters of networks rather than cleanly delineated modules. Our findings support a multivariate, interactionist model of empathic concern, in which individual differences arise from stable patterns of interaction between bottom-up systems involved in affect, motor behavior and perception and top-down systems involved in appraisal and cognitive control, regardless of task demands. Furthermore, these findings constitute a strong test of this model by showing that these characteristic patterns are observable at rest. These findings also have important diagnostic implications, as they suggest an ability to assess empathic predispositions in populations that are unable to perform conventional tests of empathic function.

